# Elevated ozone reduces photosynthetic carbon gain by accelerating leaf senescence of inbred and hybrid maize in a genotype-specific manner

**DOI:** 10.1101/154104

**Authors:** Craig R. Yendrek, Gorka Erice, Christopher M. Montes, Tiago Tomaz, Crystal A. Sorgini, Patrick J. Brown, Lauren M. McIntyre, Andrew D.B. Leakey, Elizabeth A. Ainsworth

**Author notes:** current address: The Scotts Company, Marysville, OH 43041. current address: Estación Experimental del Zaidín (EEZ-CSIC), Spain. current address: Sun Pharmaceuticals Industries, Latrobe, Tasmania (Tiago Tomaz). Corresponding author: Elizabeth Ainsworth, 1201 W. Gregory Drive, 147 ERML, Urbana, IL 61801 USA,).

## Abstract

Exposure to elevated tropospheric ozone concentration ([O_3_]) accelerates leaf senescence in many C_3_ crops. However, the effects of elevated [O_3_] on C_4_ crops including maize (*Zea mays* L.) are poorly understood in terms of physiological mechanism and genetic variation in sensitivity. Using Free Air gas Concentration Enrichment (FACE), we investigated the photosynthetic response of 18 diverse maize inbred and hybrid lines to season-long exposure to elevated [O_3_] (~100 nL L^−1^) in the field. Gas exchange was measured on the leaf subtending the ear throughout the grain filling period. On average over the lifetime of the leaf, elevated [O_3_] led to reductions in photosynthetic CO_2_ assimilation of both inbred (-22%) and hybrid (-33%) genotypes. There was significant variation among both inbred and hybrid lines in the sensitivity of photosynthesis to elevated [O_3_], with some lines showing no change in photosynthesis at elevated [O_3_]. Based on analysis of inbred line B73, the reduced CO_2_ assimilation at elevated [O_3_] was associated with accelerated senescence decreasing photosynthetic capacity, and not altered stomatal limitation. These findings across diverse maize genotypes could advance the development of more ozone tolerant maize, and provide experimental data for parameterization and validation of studies modeling how O_3_ impacts crop performance.

## INTRODUCTION

Tropospheric ozone (O_3_) is an airborne pollutant that enters leaves through their stomata and generates reactive oxygen species (ROS) upon contact with intercellular leaf surfaces (Heath 1988). Subsequently, these ROS can cause oxidative damage to membranes and other cellular components. At very high concentrations, O_3_ can elicit a hypersensitive response, but even at lower concentrations O_3_ can impair cellular function (Krupa & Manning 1988). When exposed to elevated O_3_ concentrations ([O_3_]) throughout the growing season, many plants have reduced photosynthetic C assimilation and biomass production as well as increased antioxidant capacity and mitochondrial respiration rates (Fiscus *et al*. 2005; Ainsworth *et al.* 2012). As a cumulative result of these physiological responses, crops exhibit significant reductions in economic yield with increasing [O_3_] (Mills *et al.* 2007; Ainsworth 2017).

The effects of exposure to chronic, elevated [O_3_] on photosynthesis and stomatal conductance can vary with leaf age and canopy position (Tjoelker *et al*. 1995; Morgan *et al*. 2004; Betzelberger *et al*. 2010; Feng *et al*. 2011). Studies that repeatedly measured a cohort of soybean and wheat leaves found that photosynthesis and stomatal conductance were not significantly affected by elevated [O_3_] in recently mature leaves, but continued exposure over time accelerated the decline in photosynthetic capacity as leaves aged (Morgan *et al*. 2004; Feng *et al*. 2011; Emberson *et al*., 2017). Stomatal conductance can also become uncoupled from photosynthesis as leaves age under O_3_ stress, often with greater measured reductions in photosynthesis than stomatal conductance (Lombardozzi *et al*. 2012), resulting in decreased water use efficiency at elevated [O_3_] (Tjoelker *et al*. 1995; VanLoocke *et al*. 2012). Additionally, stomata of plants exposed to elevated [O_3_] can show delayed opening or closing responses to environmental or hormonal signals (Keller & Häsler 1984; Wilkinson & Davies 2009; Paoletti & Grulke 2010; Wagg *et al*. 2013).

Much of our understanding of the effects of elevated [O_3_] on photosynthesis and stomatal function come from study of C_3_ species (Reich 1987; Paoletti & Grulke 2005; Felzer *et al*. 2007; Wittig *et al*. 2007). A few experiments have shown that elevated [O_3_] can negatively impact photosynthetic properties and biomass production of the model C_4_ species, maize (*Zea mays* L.) (Pino *et al*. 1995; Leitao *et al.* 2007a; Leitao *et al.* 2007b; Bagard *et al*. 2015; Yendrek *et al*. 2017). However, current understanding of the mechanisms underlying photosynthetic responses to O_3_ in maize is limited to a few genotypes, and often to juvenile growth stages. The high ratio of photosynthetic CO_2_ assimilation relative to stomatal conductance and isolation of Rubisco carboxylation in the bundle sheath cells make it possible that C_4_ photosynthesis will respond distinctly to C_3_ photosynthesis at elevated [O_3_]. Analysis of historical U.S. maize yields suggests that reported physiological responses to O_3_ pollution drive annual yield reductions of approximately 10% from 1980-2011, amounting to $7.2 billion in lost profit per year on average (McGrath *et al.* 2015). Considering that O_3_-mediated yield reductions were more prominent in hot and dry years (McGrath *et al.* 2015) and future climate models predict an increase in growing season temperature with altered frequency and magnitude of precipitation events (Christensen *et al.* 2007; Walsh *et al.* 2014), efforts to address the knowledge gap about mechanisms of O_3_-sensitivity in maize are likely to become increasingly important.

In addition to being the most widely produced food crop, maize is a model C_4_ species with considerable resources for analysis of genotype to phenotype associations. This includes the maize nested association mapping (NAM) population, which was developed to encompass much of the existing diversity in a relatively small number of inbred lines (Yu *et al.* 2008). Substantial phenotypic diversity for many agronomic traits exists within the parental lines of the NAM (Flint-Garcia *et al.* 2005). In order to exploit these resources to locate genetic factors associated with complex traits such as O_3_ tolerance, there is a need to first identify lines that respond differently to elevated [O_3_] for key physiological traits. For understanding and modeling the impacts of O_3_ on crops, photosynthesis and stomatal conductance are two key traits (Reich 1987; Emberson *et al*. 2000; Sitch *et al*. 2007). In this study, we investigated the effects of season-long growth at elevated [O_3_] on maize photosynthetic gas exchange. Maize was exposed to elevated [O_3_] in the field using Free Air gas Concentration Enrichment (FACE). In 2013 and 2014, we studied the inbred cultivar B73. We hypothesized that growth at elevated [O_3_] would reduce photosynthetic capacity and daily C gain as leaves progressed more rapidly through senescence. In 2015, we investigated the response of photosynthesis and stomatal conductance to elevated [O_3_] in 10 diverse inbred and 8 diverse hybrid lines in order to test for genetic variation in O_3_ response.

## MATERIALS AND METHODS

### Field site and experimental conditions

Maize (*Zea mays*) inbred and hybrid lines (Table 1) were studied at the FACE research facility in Champaign, IL (<u>www.igb.illinois.edu/soyface/</u>) in 2013, 2014 and 2015. This facility is located on a 32 ha farm. Maize is grown on half of the land each year, and rotated annually with soybean. Each year, the portion of the field growing maize was fertilized with N (180 lbs per acre) and treated with pre-emergent and post-emergent herbicides at the recommended rates. Additional weeding was done inside the FACE rings as needed. Maize inbred and hybrid lines were planted with a precision planter in rows spaced 0.76 m apart and 3.35 m in length, at a density of 8 plants m^−1^ in replicated blocks (n=4). Each block had one ambient plot and one elevated [O_3_] plot, which were octagonal in shape and of 20-m diameter. Weather conditions were monitored at the FACE facility, and Supplemental Figure 1 shows the seasonal time courses of: (1) daily maximum, (2) daily minimum temperatures, and (3) daily total precipitation for the 2013, 2014 and 2015 growing seasons.

**Table 1.** Planting date (day of year), inbred and hybrid lines examined in each field season, and the date that anthesis was recorded for each line.

| Year | Planting<br>(DOY) | Line | Anthesis<br>(DOY) |
| --- | --- | --- | --- |
| 2013 | 136, 137 | B73 | 208 |
| 2014 | 139 | B73 | 210 |
| 2015 | 139 | B73 | 205 |
|  |  | C123 | 203 |
|  |  | CML333 | 214 |
|  |  | Hp301 | 205 |
|  |  | IL14H | 201 |
|  |  | M37W | 211 |
|  |  | Mo17 | 204 |
|  |  | MS71 | 202 |
|  |  | NC338 | 214 |
|  |  | OH43 | 201 |
|  |  | B73xC123 | 202 |
|  |  | B73xCML333 | 207 |
|  |  | B73xHP301 | 202 |
|  |  | B73xIL14H | 201 |
|  |  | B73xM37W | 206 |
|  |  | B73xMo17 | 202 |
|  |  | B73xMS71 | 201 |
|  |  | B73xOH43 | 201 |

Air enriched with O_3_ was delivered to the experimental rings with FACE technology, as described in Yendrek *et al*. (2017). The target fumigation set-point in the elevated [O_3_] rings was 100 nL L^−1^, which was imposed for ~8 h per d throughout the growing season except when leaves were wet or wind speeds were too low to ensure accurate fumigation (i.e., <0.5 m s^−1^). Over the three years, when the fumigation was on, the averaged 1-min [O_3_] within the treatment rings was within 10% of the 100 nL L^−1^ set point for 53.7% of the time and within 20% of the set point for 78% of the time. Season-long 8 hr average [O_3_] in ambient rings was 40.8 nL L^−1^ in 2013, 40.1 nL L^−1^ in 2014, and 40.0 nL L^−1^ in 2015, and season-long elevated [O_3_] was 71.0 nL L^−1^ in 2013, 70.8 nL L^−1^ in 2014 and 64.0 nL L^−1^ in 2015.

### Leaf-level gas exchange measurements

*In situ* photosynthetic gas exchange of the leaf subtending the ear was measured using portable photosynthesis systems (LI-6400, LICOR Biosciences, Lincoln, NE) in a modified version of a previously published protocol (Leakey *et al.* 2006). The net rate of photosynthetic CO_2_ assimilation (*A*), stomatal conductance (*g_s_*), the ratio of intercellular [CO_2_] (c_i_) to atmospheric [CO_2_] (c_a_), and instantaneous water use efficiency (iWUE = A/g_s_) were measured. Reproductive development (time to silking and anthesis) was monitored in all experimental rows, and all dates of gas exchange measurements are reported relative to the date of anthesis.

In 2013 and 2014, the diurnal time course of leaf photosynthetic gas exchange was assessed with measurements taken across all replicate plots every few hours throughout the day at three or four development stages over the growing season of the B73 inbred genotype. In 2015, *in situ* gas exchange measurements on 10 inbred and 8 hybrid lines were collected at midday approximately every 7 d starting when the plants reached anthesis and ending when the ear leaf of >50% plants in an individual row had fully senesced. After senescence of a genotype in a given treatment, we recorded zero for *A* and g_s_ for all subsequent weekly measurements until both ambient and elevated plots were fully senesced. In all years, two or three plants were measured as sub-samples within each replicate plot at either ambient [O_3_] or elevated [O_3_].

Prior to a set of measurements being performed over a 1-2 h period of a day in all replicate plots, temperature and relative humidity (RH) in the leaf chamber cuvette were set to match prevailing ambient conditions as measured by an on-site weather station. A linear quantum sensor (AccuPAR LP-80; Decagon Devices, Pullman, WA) was used to measure the average light intensity at the position in the canopy for the leaf subtending the ear, which was used to set the PPFD incident on the leaf in the gas exchange cuvette. In 2015, a constant PPFD was used for all weekly midday measurements based on the measured PPFD at anthesis. For inbred cultivars, PPFD was set at 1,800 μmol m^−2^ s^−1^ and for hybrid cultivars PPFD was set at 450 μmol m^−2^ s^−1^. Keeping light constant over the entire measurement period enabled comparison of changes in the response of *A* and *g*_s_ to leaf aging in ambient and elevated [O_3_], and eliminated variation in those parameters due to PPFD. On all dates, four instruments were used simultaneously. All environmental conditions were held constant across instruments.

The response of *A* to c_i_ was measured on the leaf subtending the ear in 2013 and 2014 to examine the effects of elevated [O_3_] on the maximum apparent rate of phosphoenolpyruvate carboxylase activity (*V*_pmax_) and on CO_2_-saturated photosynthetic rate (*V*_max_). Two to three leaves per ring were measured. Leaves were excised pre-dawn, immediately re-cut under water, and measured in a field laboratory. This approach minimized short-term decreases in water potential, chloroplast inorganic phosphate concentration, and maximum PSII efficiency that can occur in the field, and also enabled more leaves to be measured on multiple portable photosynthesis systems over a short period of time (Leakey *et al*. 2006). Measurements were initiated at ambient c_i_, then the reference [CO_2_] in the leaf cuvette was stepped down to 25μmol mol^−1^, before it was increased stepwise to 1200 μmol mol^−1^ while keeping PPFD constant at 2,000 μmol m^−2^ s^−1^. The initial slope of the *A*/c_i_ curve (c_i_ < 60 μmol mol^−1^) was used to calculate *V*_pmax_ according to von Caemmerer (2000). A four parameter nonrectangular hyperbolic function was used to estimate *V*max as the horizontal asymptote of the *A*/c_i_ curve. Stomatal limitation to *A* (*l*) was estimated from *A*/c*i* curves using the approach described by Farquhar and Sharkey (1982), and modified by Markelz *et al*. (2011). Mean values of *V*_pmax_ and *V*_max_ from *A*/c_i_ curves were used in combination with *in situ* values of c_i_ measured at midday to estimate the range of *l* and mean *l* for recently mature leaves (DOY 219 in 2013, DOY 213 in 2014) and senescing leaves (DOY 238 in 2013, DOY 245 in 2014) in ambient and elevated [O_3_].

### Statistical analysis

Diurnal gas exchange parameters (*A*, *g_s_*) measured in 2013 and 2014 were analyzed with a repeated mixed model ANOVA with the Kenwood-Rogers option and compound symmetry covariance structure (SAS, version 9.4, Cary, NC). All analysis was done on the ring mean. Days were analyzed independently, time of day was a repeated measure and block was a random effect. A mixed model ANOVA with the Kenwood-Rogers specification was used to analyze *V*_pmax_ and *V*_max_ in 2013 and 2014 (SAS, Version 9.4, Cary, NC). O_3_ treatment and DOY were fixed effects in the model, block was a random effect, and years were analyzed independently. Statistical differences in least squared mean estimates were determined by linear contrasts with a threshold *p*<0.05.

*A* and g_s_ of the flag leaf measured in 2015 were fit with a quadratic equation: y = y_o_ + αx +βx^2^, where y was *A* or g_s_, and x was days after anthesis (Proc Nlin, SAS). Other models (exponential decay, 3-parameter Weibull function) were tested, but did not fit all of the data for each genotype. Therefore, a simple quadratic equation was used. In order to test if there were differences in the decline in *A* or g_s_ over time in ambient and elevated [O_3_], a single quadratic model was first fit to the *A* and g_s_ data for each genotype, then models were fit to each genotype and treatment combination. An F statistic was used to test if the model with genotype and treatment produced a significantly better fit to the data, i.e., if there were significant differences in the response of *A* or g_s_ over time in ambient and elevated [O_3_]. Additionally the quadratic model was fit to each genotype within each ring in order to obtain parameter estimates for y_o_, α and β. Parameter estimates as well as initial values of *A* and g_s_ measured shortly after anthesis were analyzed with a mixed model ANOVA with the Kenwood-Rogers specification. O_3_ treatment and genotype were fixed effects in the model and block was a random effect. Statistical differences in least squared mean estimates between ambient and elevated [O_3_] for each inbred or hybrid line were determined by linear contrasts with a threshold *p*<0.05.

## RESULTS

### Elevated [O_3_] accelerates loss of photosynthetic capacity in inbred line B73

In 2013 and 2014, daily time courses of leaf photosynthetic gas exchange were measured *in situ* as the leaf subtending the ear aged in ambient and elevated [O_3_] (Figs. 1, 2). When the leaf was recently fully expanded, shortly before or after anthesis, there was no significant effect of elevated [O_3_] on *A* (Figs. 1d, 2e, 2f). However, as the leaf aged, growth at elevated [O_3_] accelerated the decline in diurnal C gain, and later in the season *A* was significantly reduced by elevated [O_3_] (Figs. 1e, 1f, 2g, 2h). Significant differences in *A* between ambient and elevated [O_3_] were apparent earlier in the growing season than differences in g_s_ (Figs. 1e versus 1h, 2g versus 2k). But, there were no consistent differences in the ratio of intercellular [CO_2_] to atmospheric [CO_2_] (c_i_/c_a_) or instantaneous water use efficiency (iWUE = *A*/g_s_) in ambient and elevated [O_3_] in either year (Fig. 1j-o, Fig. 2m-t). On only one sampling date out of seven, which was characterized by low PPFD, elevated [O_3_] led to significantly greater c_i/_c_a_ and significantly lower iWUE (Fig. 1k, n).

**Figure 1.**
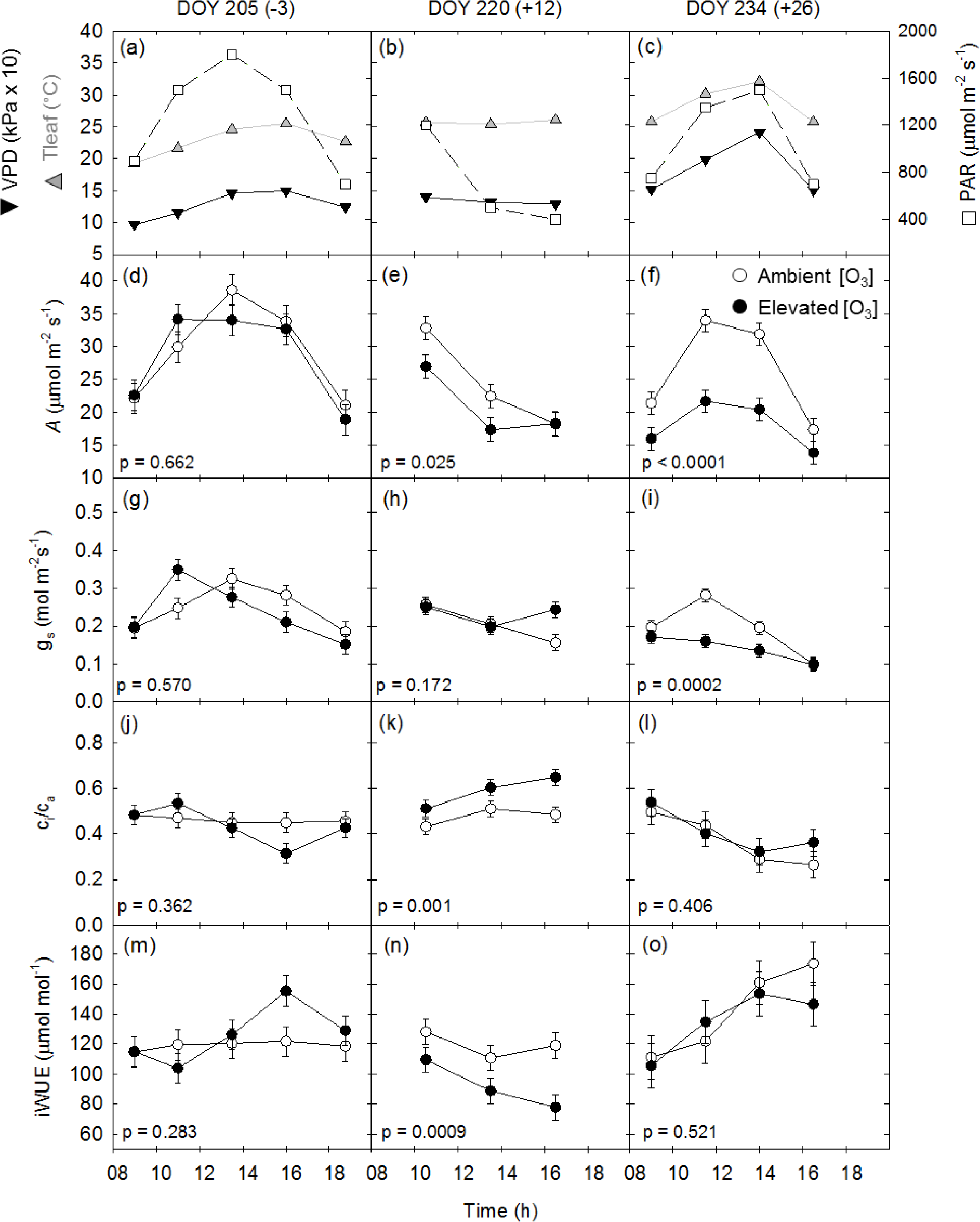
The diurnal response of net assimilation (*A*) and stomatal conductance (*g*_s_) of maize inbred line B73 in 2013. Measurements were made on the leaf subtending the ear on three days of the year (DOY). The number of days from anthesis is shown in parenthesis after measurement DOY. The conditions in the leaf cuvette including leaf temperature (T_leaf_), vapor pressure deficit (VPD), and photosynthetically active radiation (PAR) are given for each date in panels a, b and c. Diurnal *A* is shown for each DOY in panels d, e and f, and *g*_s_ is shown in panels g, h and i. The least squared means ± 1 standard error are shown. The *p*-value represents the O_3_ effect for each DOY.

**Figure 2.**
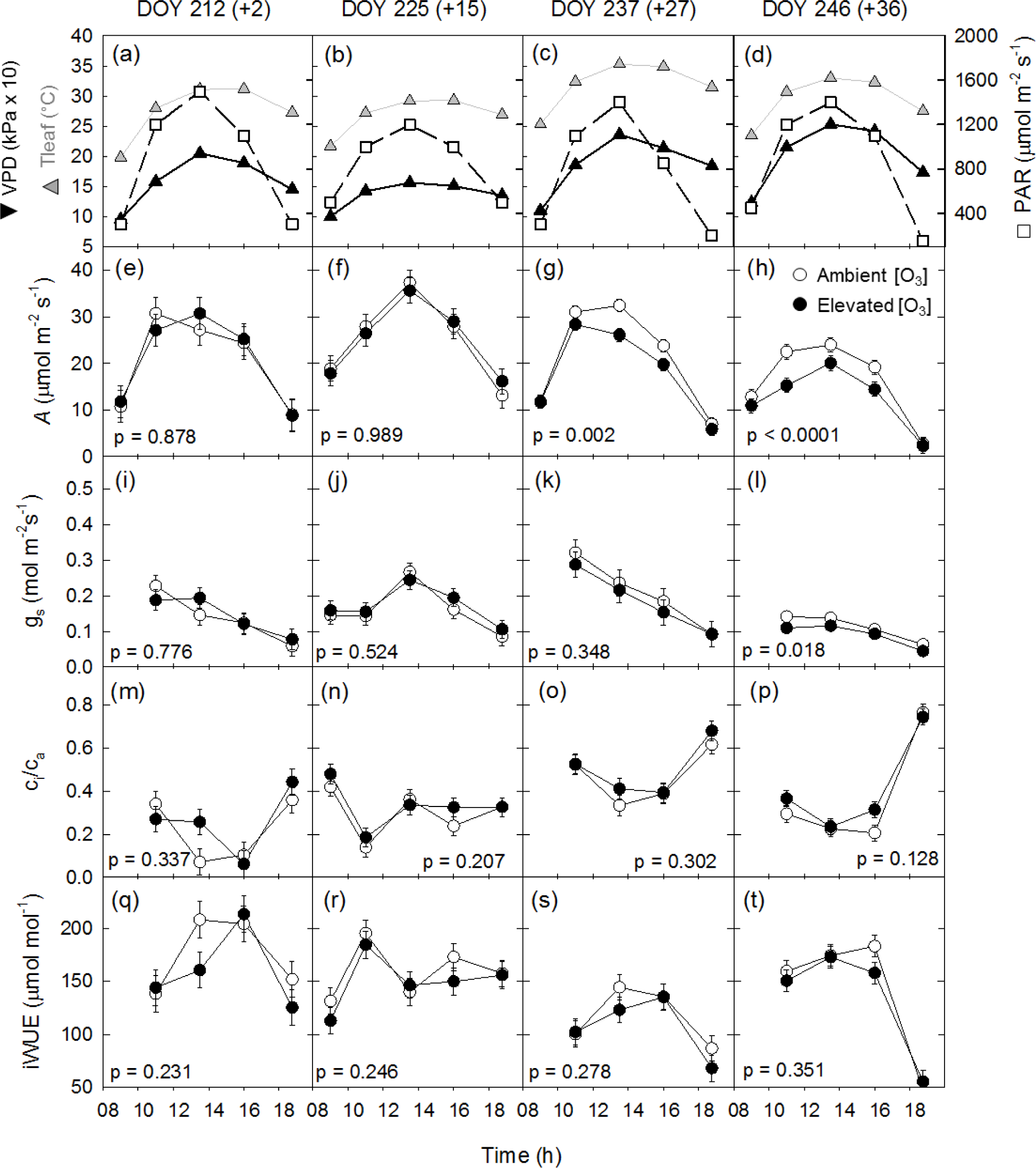
The diurnal response of net assimilation (*A*) and stomatal conductance (*g*_s_) of maize inbred line B73 in 2014. Measurements were made on the leaf subtending the ear on four days of the year (DOY). The number of days from anthesis is shown in parenthesis after measurement DOY. The conditions in the leaf cuvette including leaf temperature (T_leaf_), vapor pressure deficit (VPD), and photosynthetically active radiation (PAR) are given for each date in panels a, b, c and d. Diurnal *A* is shown for each DOY in panels e, f, g and h, and *g*_s_ is shown in panels i, j, k and l. The least squared means ± 1 standard error are shown. The *p*-value represents the O_3_ effect for each DOY.

Declines in *in situ A* over the leaf aging process were associated with decreased photosynthetic capacity, measured as *V*_pmax_ (Fig. 3a, b; Table 2) and *V*_max_ (Fig. 3c, d; Table 2). Declines in *V*_pmax_ from when the leaf was recently fully expanded to until ~30 days later were greater in elevated [O_3_] (-64% in 2013 and −64% in 2014) than in ambient [O_3_] (-51% in 2013 and −33% in 2014; Fig. 3a, b). Likewise, declines in *V*_max_ from when the leaf was recently fully expanded to until ~30days later were greater in elevated [O_3_] (-40% in 2013 and −52% in 2014) than in ambient [O_3_] (-25% in 2013 and −33% in 2014; Fig. 3c, d). Although an O_3_ treatment effect was not significant in the mixed ANOVA model (Table 2), pair-wise comparisons of the means showed that *V*_pmax_ was significantly lower in elevated [O_3_] on the final sampling date in 2014 (Fig. 3b) and *V*_max_ was significantly lower in elevated [O_3_] on the final sampling date in both 2013 and 2014 (Fig. 3c, d).

**Figure 3.**
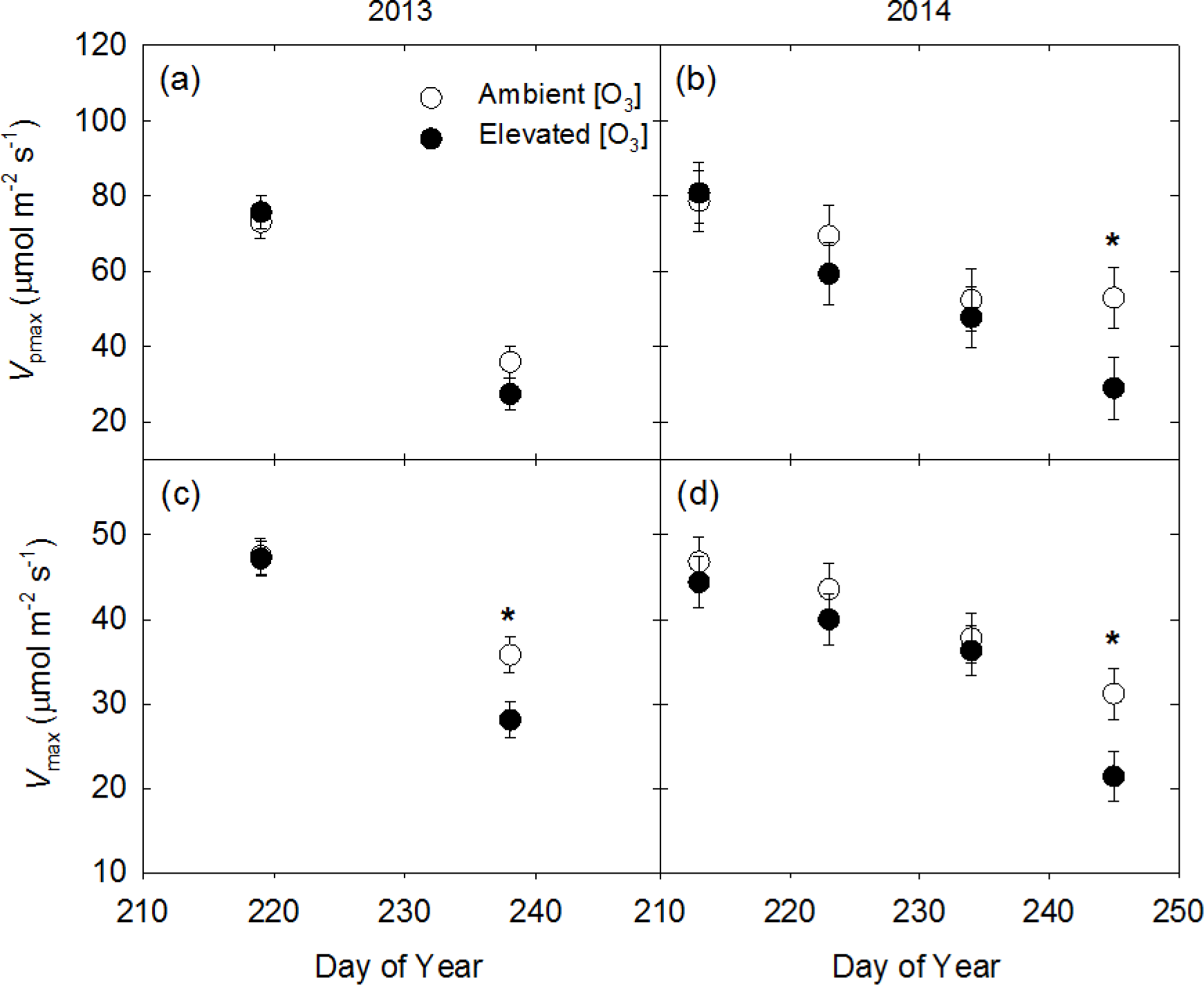
Maximum carboxylation capacity of PEPC (*V*_pmax_) (a, b) and CO_2_-saturated photosynthetic rate (*V*_max_) (c, d) of maize inbred line B73 measured in 2013 (a, c) and 2014 (b, d). Measurements were made on the leaf subtending the ear. The least squared means ± 1 standard error are shown. Significant differences between ambient and elevated [O_3_] (*p*<0.05) for individual days are indicated with an asterisk.

**Table 2.** Analysis of variance (*F*, p) of the maximum apparent rate of phosphoenolpyruvate carboxylase activity (*V*_pmax_) and CO_2_- saturated photosynthetic rate (*V*_max_) measured in inbred line B73 grown at ambient and elevated [O_3_] in 2013 and 2014. Years were analyzed independently.

|  |  | 2013 |  | 2014 |  |
| --- | --- | --- | --- | --- | --- |
| | | $F$ | $p$ | $F$ | $p$ |
| $V_{\text{pmax}}$ | $\text{O}_3$ | 0.39 | 0.5754 | 2.46 | 0.1296 |
|  | DOY | 129 | <0.0001 | 8.57 | 0.0005 |
| | $\text{O}_3 \times \text{DOY}$ | 2.22 | 0.1867 | 0.92 | 0.4457 |
| $V_{\text{max}}$ | $\text{O}_3$ | 2.97 | 0.1358 | 3.02 | 0.1327 |
|  | DOY | 67.9 | 0.0002 | 18.3 | <0.0001 |
| | $\text{O}_3 \times \text{DOY}$ | 4.01 | 0.0921 | 0.94 | 0.4411 |

*A*/c_i_ curves were also used to assess stomatal vs. biochemical limitations to *A* in aging leaves grown at ambient and elevated [O_3_]. Average *A*/c_i_ curves in ambient and elevated [O_3_] were plotted for the leaf subtending the ear shortly after anthesis, and then ~3-4 weeks later (Fig. 4). Shortly after anthesis, *l* was minimal in both ambient and elevated [O_3_] in 2013 (2-3%), and mean c_i_ was well above the inflection point of the *A*/c_i_ curve (Fig. 4a). In 2014, there was a wider range of observed c_i_ shortly after anthesis, but mean *l* was still low (5-6%) in both ambient and elevated [O_3_], and the mean c_i_ was above the inflection point (Fig 4b). As leaves aged, *l* increased in both ambient and elevated [O_3_] to 20-30% (Fig. 4c, d). Both increased *l* and decreased photosynthetic capacity result in lower *A* in aging leaves, but elevated [O_3_] did not appear to change *l*.

**Figure 4.**
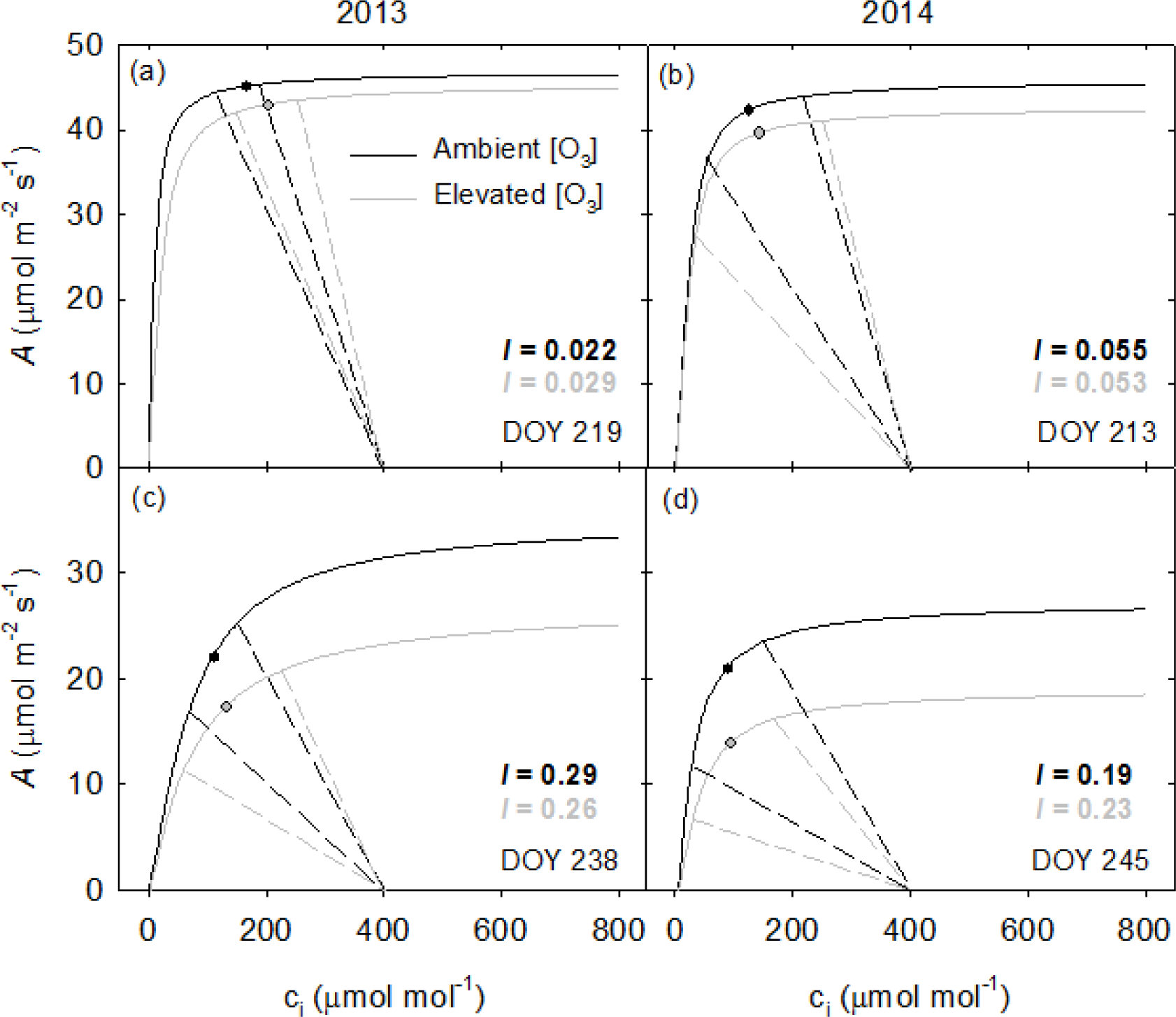
Summary of *A*/c_i_ response curves (solid lines) and CO_2_ supply functions (dashed lines) for maize inbred line B73 grown at ambient [O_3_] (black lines) and elevated [O_3_] (grey lines). The supply functions show the maximum and minimum c_i_ observed at midday in the field during diurnal measurements of *A*, and the points on the curves show the mean observed ci. Measurements were made on the leaf subtending the ear shortly after anthesis (a, b) and ~4-5 weeks after anthesis (c, d). Estimates of mean stomatal limitation (*l*) are reported in each panel for ambient and elevated [O_3_].

### Photosynthetic response to elevated [O_3_] in diverse inbred and hybrid lines

In 2015, *A* of the leaf subtending the ear was measured approximately weekly in 10 diverse inbred lines and 8 hybrid lines during the grain filling period. Initial values of *A* measured shortly after anthesis differed among lines, but not between ambient and elevated [O_3_], and the interaction between inbred line and [O_3_] was only marginally significant (Table 3). Similarly, initial measurements of g_s_ varied among inbred lines, but not between ambient and elevated [O_3_] (Table 3). For 8 of the 10 inbred lines, there was significant acceleration in the decline in *A* as the leaf aged under elevated [O_3_], similar to what was observed in previous years in B73 (Fig. 5). However, for two inbred lines (M37W and CML333), there was no significant difference in the quadratic fit to the photosynthetic data between ambient and elevated [O_3_] (Fig. 5). There were highly significant effects of [O_3_], inbred line and their interaction on the parameters describing the quadratic equation modeling the decline in *A* over time in the inbred lines (Table 3). This reflected variation in the estimates of *A* at anthesis (*y*_o_), the time to the start of a decline in *A* (α) and the rate of decline in *A* (β). The significant effect of [O_3_] on *y*_o_ is not consistent with the lack of an effect on initial rates of *A*, but some of the data were best fit by convex functions and some by concave functions which affected accurate prediction of *y*_o_ in the inbred lines (Fig. 5). Average α was 0.30 in ambient [O_3_] and −0.69 in elevated [O_3_], indicating that a decline in *A* started earlier in elevated [O_3_] (Fig. 5, Supplemental Table 1). Average β was −0.016 in ambient [O_3_] and −0.00025 in elevated [O_3_] indicating that the rate of decline in *A* was greater in ambient [O_3_] than elevated [O_3_] on average across inbred lines (Supplemental Table 1).

**Table 3.** Analysis of variance (*F*, p) of photosynthesis (*A*), stomatal conductance (g_s_) and the quadratic parameters describing the decline in *A* and g_s_ over time. Measurements were made in in 10 inbred and 8 hybrid lines grown at ambient and elevated [O_3_] in 2015, and inbreds and hybrids were analyzed in separate models. *A* and g_s_ represent the first measurement taken shortly after anthesis in Figs. 5-8.

| **Inbred Lines** |  |  | ***A*** |  |  | ***g<sub>s</sub>*** |  |  |
| --- | --- | --- | --- | --- | --- | --- | --- | --- |
|  | *A* (μmol m<sup>-2</sup> s<sup>-1</sup>) | *g<sub>s</sub>* (mol m<sup>-2</sup> s<sup>-1</sup>) | *y*<sub>0</sub> | α | β | *y*<sub>0</sub> | α | β |
| [*O*<sub>3</sub>] | 1.33, 0.254 | 0.26, 0.609 | 13.1, 0.0006 | 28.8, <0.0001 | 28.0, <0.0001 | 2.95, 0.091 | 6.92, 0.011 | 7.65, 0.007 |
| Line | 2.93, 0.006 | 2.89, 0.007 | 12.9, <0.0001 | 13.0, <0.0001 | 9.19, <0.0001 | 8.19, <0.0001 | 7.67, <0.0001 | 5.96, <0.0001 |
| [*O*<sub>3</sub>] x Line | 1.87, 0.076 | 1.77, 0.094 | 3.12, 0.004 | 5.51, <0.0001 | 5.76, <0.0001 | 1.23, 0.296 | 2.29, 0.028 | 2.58, 0.014 |
| **Hybrid Lines** |  |  | ***A*** |  |  | ***g<sub>s</sub>*** |  |  |
|  | *A* (μmol m<sup>-2</sup> s<sup>-1</sup>) | *g<sub>s</sub>* (mol m<sup>-2</sup> s<sup>-1</sup>) | *y*<sub>0</sub> | α | β | *y*<sub>0</sub> | α | β |
| [*O*<sub>3</sub>] | 0.00, 0.965 | 0.00, 0.953 | 55.9, 0.017 | 66.4, 0.015 | 61.8, 0.016 | 1.82, 0.310 | 4.15, 0.179 | 4.72, 0.095 |
| Line | 1.04, 0.429 | 0.39, 0.902 | 1.52, 0.202 | 1.67, 0.157 | 4.50, 0.002 | 0.77, 0.618 | 1.80, 0.128 | 3.98, 0.004 |
| [*O*<sub>3</sub>] x Line | 0.58, 0.765 | 1.19, 0.340 | 5.72, 0.0004 | 8.16, <0.0001 | 10.4, <0.0001 | 1.60, 0.177 | 1.97, 0.095 | 2.36, 0.050 |

**Figure 5.**
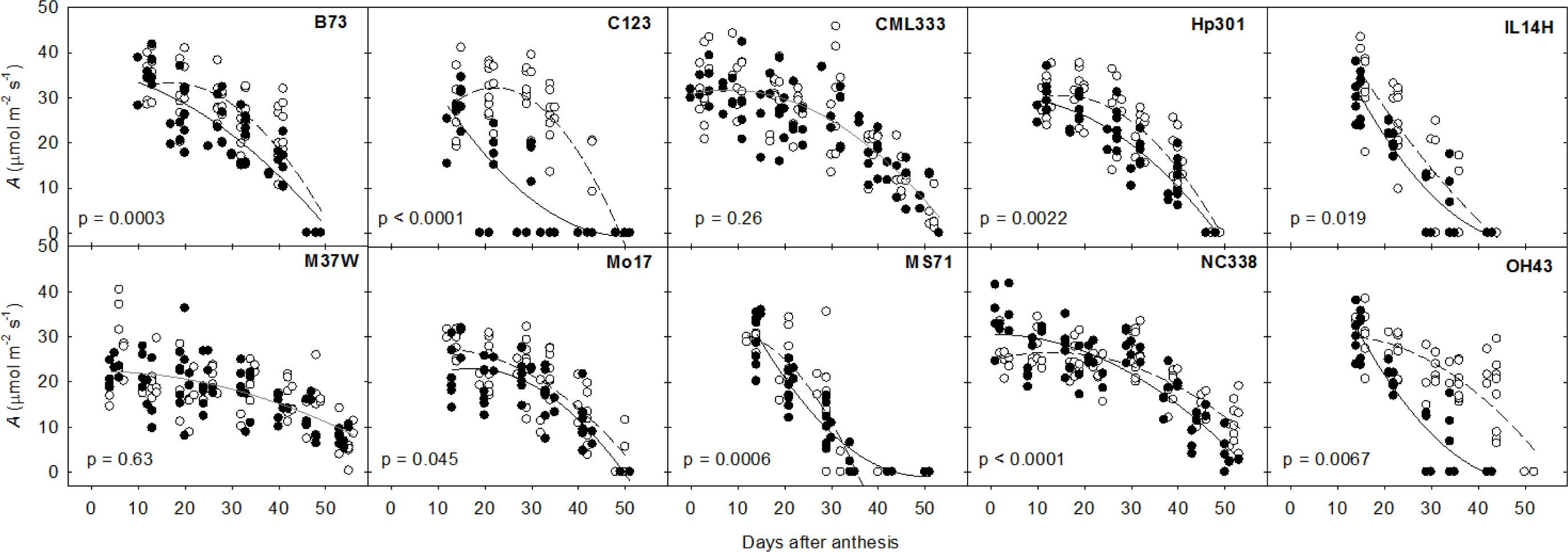
Photosynthetic rate (A) of the leaf subtending the ear measured during grain filling in 2015 for 10 inbred lines grown at ambient (open circles) and elevated [O_3_] (filled circles). The best fit quadratic curves for ambient (dashed line) and elevated [O_3_] (solid line) are shown if the statistical test revealed that there was a better fit to the data using two quadratic curves (i.e., one for each treatment) rather than a single curve for all of the data. Grey lines represent the best quadratic fit through all of the data when the statistical test revealed no difference between ambient and elevated [O_3_]. Each point represents the mean value for an individual replicate ring for each measurement date. Parameters describing these curves (*y*_o_, α, β) are reported in Supplemental Table 1.

Measurements of g_s_ in the aging flag leaf in inbred lines differed from the response of *A* over time. For 7 of the 10 genotypes, there was no evidence for accelerated decline in g_s_ over time (Fig. 6). Of the remaining 3 genotypes, MS71 and C123 displayed more rapid decline in g_s_ over time, while NC338 had initially higher g_s_ in elevated [O_3_]. The influence of this small set of genotypes meant that there were significant effects of [O_3_], inbred line and their interaction on the shape of the quadratic fit to the decline in g_s_ over time, especially on parameters α and β (Table 3; Supplemental Table 1). Apparent differences in the response of *A* and g_s_ to elevated [O_3_] over the grain filling period did not impact the iWUE, which was not different in ambient and elevated [O_3_] (Supplemental Fig. 2).

**Figure 6.**
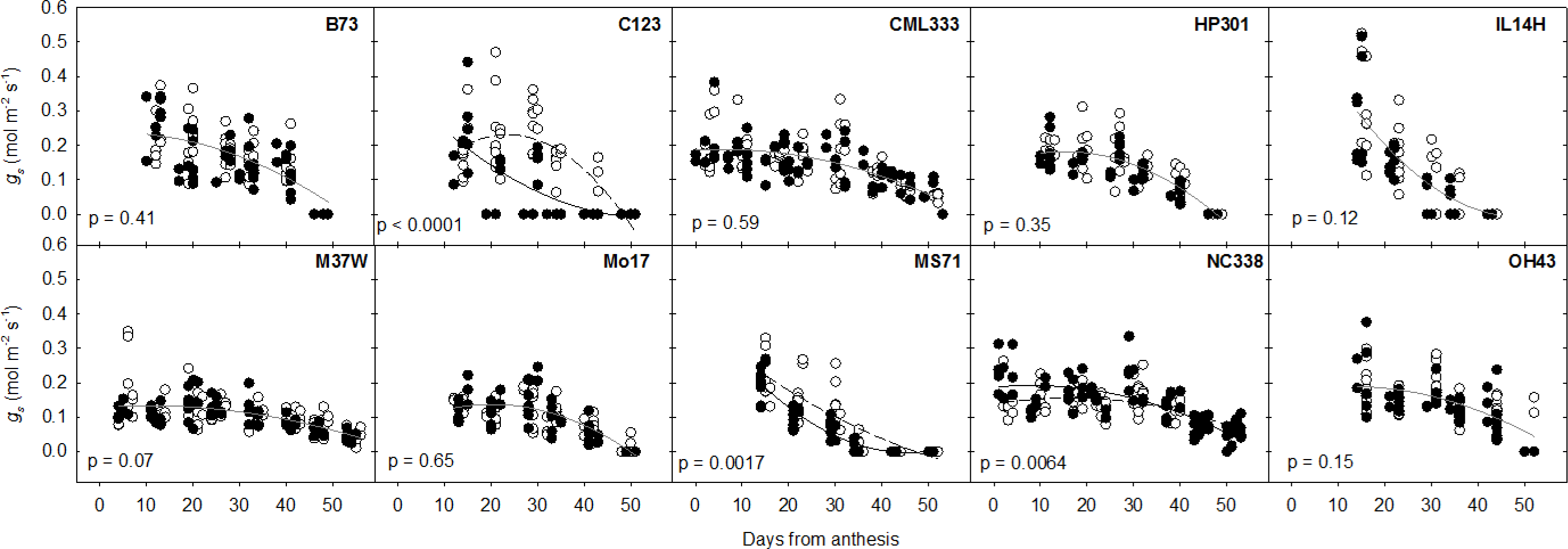
Stomatal conductance (g_s_) of the leaf subtending the ear measured during grain filling in 2015 for 10 inbred lines grown at ambient (open circles) and elevated [ O_3_] (filled circles). The best fit quadratic curves for ambient (dashed line) and elevated [O_3_] (solid line) are shown if the statistical test revealed that there was a better fit to the data using two quadratic curves (i.e., one for each treatment) rather than a single curve for all of the data. Grey lines represent the best quadratic fit through all of the data when the statistical test revealed no difference between ambient and elevated [O_3_]. Each point represents the mean value for an individual replicate ring for each measurement date. Parameters describing these curves (y_0_, α, β) are reported in Supplemental Table 1.

All 8 of the hybrid lines showed acceleration in the decline of *A* in the leaf subtending the ear over the grain filling period (Fig. 7). Unlike the inbreds, the hybrids had similar initial values of *A* and g_s_ shortly after anthesis (Table 3). These measurements were taken at a lower light level than the inbred measurements, which was representative of the light measured in the canopy at the leaf subtending the ear in hybrids vs. inbreds, but may explain why there was less variation among hybrid lines. There was a significant effect of [O_3_] and a significant [O_3_] x hybrid line interaction on the parameters describing the decline in *A* over time (Table 3; Supplemental Table 1). Consistent with the inbred lines, α was positive in ambient [O_3_] (1.06) and negative in elevated [O_3_] (-0.033), indicating a more immediate decline in *A* in elevated [O_3_], and β was more negative in ambient [O_3_] (-0.02) than elevated [O_3_] (-0.006) indicating more rapid decline in *A* in ambient [O_3_] (Supplemental Table 1). Only 3 of the 8 hybrid lines showed significant changes in g_s_ at elevated [O_3_] over the grain filling period (Fig. 8), and consistent with the inbred lines, the apparent differences in the response of *A* and g_s_ to elevated [O_3_] over time did not impact iWUE (Supplemental Fig. 3). Notably, the hybrids of MS71 and C123 crossed with B73 were among this sensitive group, matching the greater than average sensitivity of these genotypes as inbreds.

**Figure 7.**
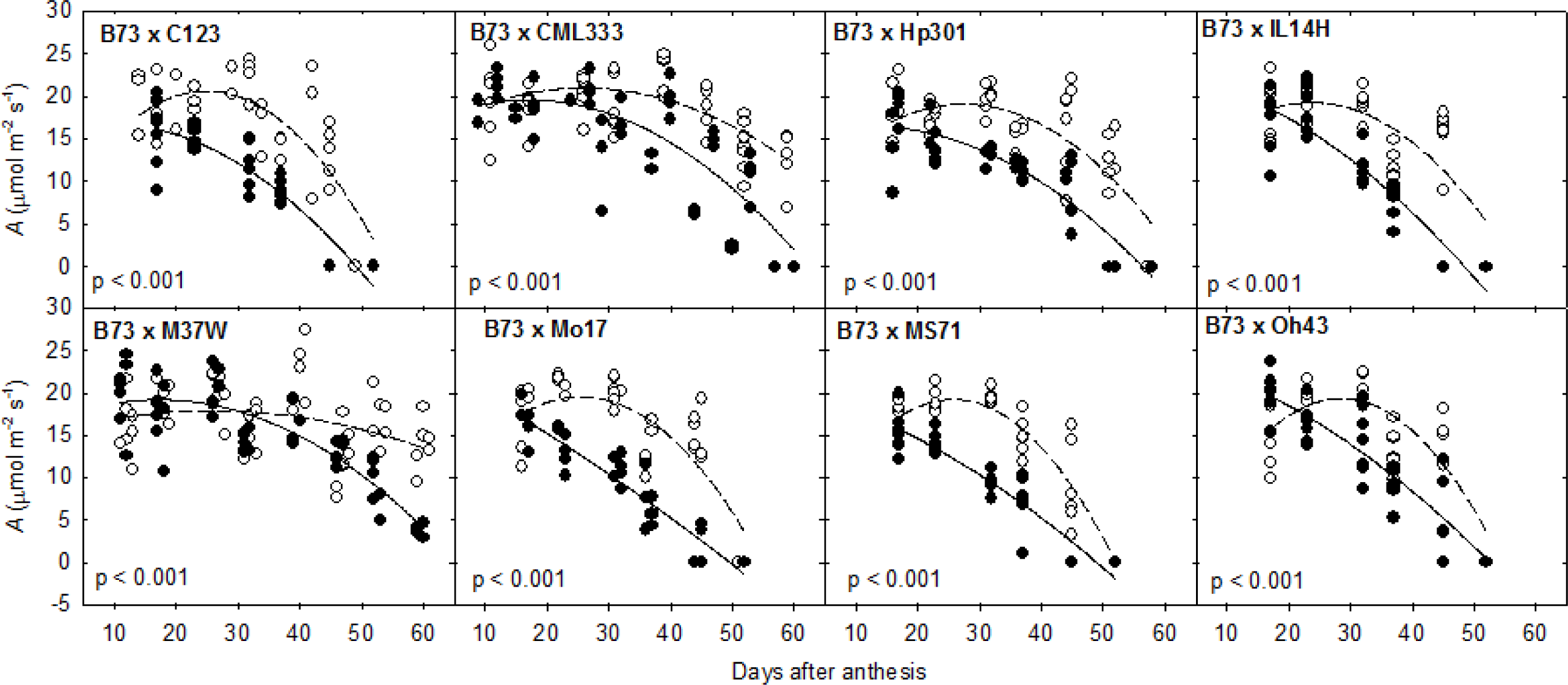
Photosynthetic rate (*A*) of the leaf subtending the ear measured during grain filling in 2015 for 8 hybrid lines grown at ambient (open circles) and elevated [O_3_] (filled circles). The best fit quadratic curves for ambient (dashed line) and elevated [O_3_] (solid line) are shown if the statistical test revealed that there was a better fit to the data using two quadratic curves (i.e., one for each treatment) rather than a single curve for all of the data. Grey lines represent the best quadratic fit through all of the data when the statistical test revealed no difference between ambient and elevated [O_3_]. Each point represents the mean value for an individual replicate ring for each measurement date. Parameters describing these curves (y_o_, α, β) are reported in Supplemental Table 1.

## DISCUSSION

Tropospheric [O_3_] continues to increase in many of the world’s crop growing regions as a result of inconsistent regulation of precursor pollutants around the globe and short- and long-distance transport of pollutants (Lin *et al*. 2017). Early studies of maize reported that it was significantly more tolerant to increasing [O_3_] than C_3_ crops like soybean or peanut (Heck *et al*. 1983). However, more recent experimental studies using open top chambers reported that maize leaf physiology is sensitive to chronic O_3_ exposure in the cultivar Chambord (Leitao *et al.* 2007a; Leitao *et al.* 2007b). Furthermore, multiple regression analysis of county-level yield data suggests that there is significant yield loss to O_3_ in the Midwest U.S. growing region (McGrath *et al*. 2015). Here, we examined how season-long fumigation with elevated [O_3_] using FACE technology in the primary area of maize production affected gas exchange of the leaf subtending the ear. We concentrated on this leaf and time period because previous research has established that most of the photosynthate used for grain filling in maize is provided by mid-canopy leaves after anthesis (Borrás *et al*. 2004), and those leaves are the last on the plant to senesce (Valentinuz & Tollenaar 2004). Diurnal measurements of B73 showed that ear leaf C assimilation was not affected by elevated [O_3_] around the time of anthesis when the leaf was recently mature; however, loss of photosynthetic capacity of that leaf was accelerated by elevated [O_3_] (Figs. 1 & 2).

The seasonal pattern of elevated [O_3_] effects on photosynthetic carbon gain over the diurnal period was reflected in measurements of *A* at midday (Figs. 1 & 2). Therefore, screening of midday *A* was used to explore genotypic variation in sensitivity to elevated [O_3_] among diverse hybrid and inbred lines of maize. On average, the sum of midday CO_2_ assimilation by the leaf subtending the ear over the period from anthesis to complete leaf senescence was reduced by 22% in inbred lines and 33% in hybrid lines (Table 4). However, sensitivity to elevated [O_3_] in inbred lines ranged from no loss in photosynthetic CO_2_ assimilation (CML333, M37W) to 59% lower CO_2_ assimilation (C123; Fig. 5, Table 4). Likewise, sensitivity to elevated [O_3_] in hybrids ranged from 13% lower photosynthetic CO_2_ assimilation (B73 x M37W) to 44% lower CO_2_ assimilation (B73 x MS71; Fig. 7, Table 4). All ozone sensitive lines showed the same developmentally dependent pattern of response as B73, with similar *A* in young leaves at ambient and elevated [O_3_], but more rapid decline in *A* over time (Fig. 5, 7). Thus, acceleration of senescence rather than decreased initial investment in photosynthetic capacity is a key response of maize to elevated [O_3_] at the leaf level. But, there is genetic variation in response that might be exploited to breed maize with improved tolerance to ozone pollution, and possibly even broader spectrum oxidative stress tolerance.

**Table 4.** The percentage change in elevated [O_3_] for the sum of midday photosynthesis (*A*) and average stomatal conductance (g_s_) of the leaf subtending the ear measured during grain filling in 2015. Data and quadratic fits for *A* and g_s_ in inbred and hybrid lines are shown in Figs. 5-8.

| **Line** | ***A*** | ***g<sub>s</sub>*** |
| --- | --- | --- |
| B73 | -21% | 0% |
| C123 | -59% | -13% |
| CML333 | 0% | 0% |
| Hp301 | -17% | 0% |
| IL14H | -34% | 0% |
| M37W | 0% | 0% |
| Mo17 | -16% | 0% |
| MS71 | -46% | -5% |
| NC338 | -2% | -12% |
| OH43 | -28% | +14% |
| B73xC123 | -39% | -31% |
| B73xCML333 | -22% | 0% |
| B73xHP301 | -32% | 0% |
| B73xIL14H | -39% | 0% |
| B73xM37W | -13% | 0% |
| B73xMo17 | -43% | 0% |
| B73xMS71 | -44% | -43% |
| B73xOH43 | -28% | -27% |

Accelerated loss of photosynthetic capacity in elevated [O_3_] has been documented in FACE experiments with soybean (Morgan *et al*. 2004; Betzelberger *et al*. 2010), wheat (Feng *et al*. 2011; 2016), rice (Pang *et al*. 2009) and now maize. In rice and wheat, accelerated senescence of the flag leaf at elevated [O_3_] shortened the grain filling duration and reduced seed yields (Gelang *et al*. 2000; Pang *et al*. 2009; Feng *et al*. 2011). Additionally, genetic variation in the response of seed yield to elevated [O_3_] was attributed in part to variation in leaf senescence (Pang *et al*. 2011; Feng *et al*. 2011) as well as to variation in detoxification and antioxidant capacity (Feng *et al*. 2010; 2016). Acceleration of senescence in the ear leaf of maize is associated with maize yield loss under a number of environmental stresses (Wolfe *et al*. 1988; Bänzinger *et al*. 1999), and greater yield potential of newer maize varieties has been associated with maintenance of photosynthetic capacity of the ear leaf for a longer period of time following anthesis (Ding *et al*. 2005). Thus, the acceleration of senescence reported here under elevated [O_3_] for both inbred and hybrid maize has the potential to contribute to grain yield losses.

We found that growth at elevated [O_3_] both accelerated the speed at which flag leaves lost photosynthetic capacity (represented by the β parameter in our quadratic model) and decreased the length of time that leaves maintained optimum photosynthetic capacity (α term in quadratic model). It would be interesting to test the response of maize lines with delayed leaf senescence or stay-green phenotypes for O_3_ response. There are a variety of genetic and physiological mechanisms that enable a stay-green phenotype (Thomas & Howarth 2000), and there may be mechanisms that are more promising under O_3_ stress than others. For example, stay-green lines with altered chlorophyll catabolism may be less O_3_-tolerant as the build-up of phytotoxic degradation products might compete with O_3_-induced reactive oxygen species for detoxification by the anti-oxidant system. However, stay-green lines with perennial tendencies or lines where senescence is initiated later, but progresses at the normal rate and with typical catabolism may have the potential to retain higher rates of photosynthesis later in the growing season, and therefore yield better in higher [O_3_].

Studies in a number of C_3_ species have indicated that biochemical limitation, and not stomatal limitation, are responsible for the sustained effects of elevated [O_3_] on *A* (Farage *et al*. 1991; Morgan *et al*. 2004; Paoletti & Grulke 2005; Feng *et al*. 2011). In C_3_ species, accelerated decline in *A* in aging leaves was associated with decreased maximum carboxylation capacity and electron transport capacity (Zheng *et al*. 2002; Morgan *et al*. 2004; Feng *et al*. 2011). We found a similar response in the maize inbred line B73 with the decline in *A* at elevated [O_3_] associated with decreased photosynthetic capacity (*V*_pmax_ and *V*_max_), and not with any change in limitation imposed by stomata. Stomatal limitation increased from 2-5% in the ear leaf when it was recently mature to 20-30% when it was 4-5 weeks older, which is consistent with the loss of stomatal control in aging leaves previously described for other species (Reich 1984). This increase in stomatal limitation occurred in both ambient and elevated [O_3_], and so there does not appear to be evidence for an uncoupling of *A* and g_s_ in elevated [O_3_] that has been described in other studies (Paoletti & Grulke 2005). Across the diverse inbred and hybrid lines grown at elevated [O_3_], both *A* and g_s_ declined with ear leaf age (Figs. 5-8), although the reduction in A was often greater than g_s_. Still iWUE remained relatively constant (Supplemental Fig. 2, 3), providing further evidence from diverse maize lines that growth at elevated [O_3_] does not result in uncoupling of *A* and g_s_.

**Figure 8.**
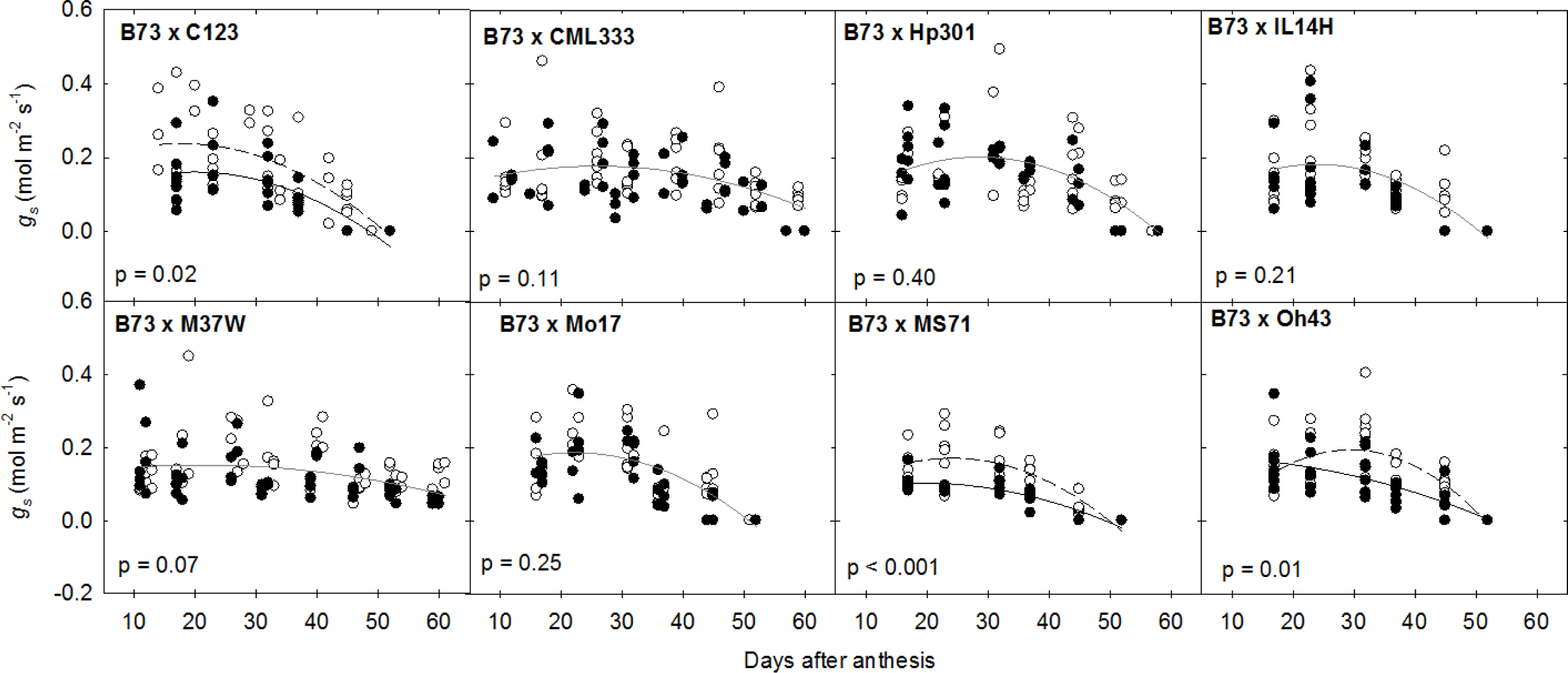
Stomatal conductance (g_s_) of the leaf subtending the ear measured during grain filling in 2015 for 8 hybrid lines grown at ambient (open circles) and elevated [O_3_] (filled circles). The best fit quadratic curves for ambient (dashed line) and elevated [O_3_] (solid line) are shown if the statistical test revealed that there was a better fit to the data using two quadratic curves (i.e., one for each treatment) rather than a single curve for all of the data. Grey lines represent the best quadratic fit through all of the data when the statistical test revealed no difference between ambient and elevated [O_3_]. Each point represents the mean value for an individual replicate ring for each measurement date. Parameters describing these curves (y_o_, α, β) are reported in Supplemental Table 1.

In addition to testing for cultivar variation in response to elevated [O_3_], we were also able to identify significant variation in *A* and g_s_ in recently mature flag leaves of inbred maize lines (Table 3). The hybrid lines, which all contained B73 as the female parent, did not show variation in *A* and g_s_ (Table 3). Previous studies have also reported that inbred lines show greater variation in photosynthetic traits compared to hybrid lines (Albergoni *et al*. 1983; Ahmadzadeh *et al*. 2004), but that hybrid lines maintain higher rates of *A* over a leaf lifespan compared to inbred lines (Ahmadzadeh *et al.* 2004). We found similar results in our studies of diverse maize lines, however, it should be duly noted that due to the recurrence of B73 as the female parent, there was more limited genetic diversity in the hybrids used in this study. Additionally, the light environment at the ear leaf at anthesis was different in inbred and hybrid canopies. *A* and g_s_ were monitored at higher PPFD in the inbred lines (1,800 μmol m^−2^ s^−1^) compared to the hybrid lines (450 μmol m^−2^ s^−1^), which may have also contributed to the greater variation observed amongst inbred lines.

Our study showed that accelerated loss of photosynthetic capacity was the basis for photosynthetic sensitivity to elevated [O_3_] in both inbred and hybrid maize lines. Drought and high temperature stress also negatively impact photosynthesis in maize (Bänzinger *et al*. 1999; Neiff *et al*. 2016), and those stresses often co-occur with high O_3_ events, and will likely increase in the future (Cook *et al*. 2014). Further experimental studies are needed to understand the interactions of rising [O_3_] with intensifying drought and heat stress, but our work suggests that a key trait for robust improvement of maize response to elevated [O_3_] is maintenance of photosynthetic capacity during the grain filling period. By testing a diverse panel of maize genotypes under field conditions in the world’s primary area of production, this study provides a foundation on which to investigate the genetic variation in maize oxidative stress tolerance, and possibly develop more stress tolerant germplasm. Under elevated [O_3_], both inbred and hybrid maize lines were shown to have reduced carbon gain of the leaf subtending the ear, which heavily influences yield. In contrast to many previous studies of crop responses to atmospheric change using FACE technology (e.g. Markelz *et al.* 2011; Gillespie *et al.* 2012), this study describes the average and variance in treatment effects across diverse genotypes, and so should provide experimental data that is a stronger basis for parameterization or validation of studies attempting to model regional crop performance.

## ACKNOWLEDGMENTS

This work was supported by the National Science Foundation (grant no. PGR–1238030) and the USDA ARS. We thank Don Ort for directing the SoyFACE facility, Alvaro Sanz for assistance with gas exchange measurements and Kannan Puthuval, Brad Dalsing and Chad Lantz for management of the FACE rings.

## FIGURE LEGENDS

**Supplemental Figure 1.** Seasonal maximum and minimum temperatures and precipitation in 2013, 2014 and 2015 measured at the FACE experimental site in Champaign, IL.

**Supplemental Figure 2.**
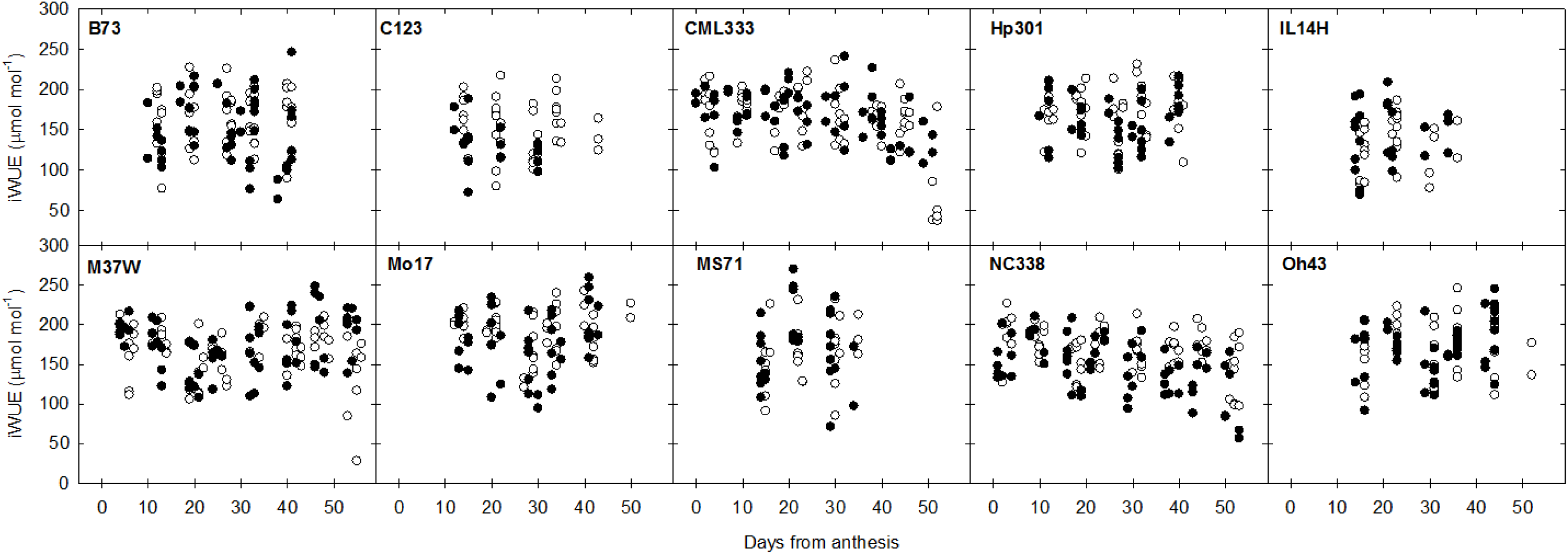
Instantaneous water use efficiency (iWUE = *A*/g_s_) of the leaf subtending the ear measured during grain filling in 2015 for 10 inbred lines grown at ambient (open circles) and elevated [O_3_] (filled circles).

**Supplemental Figure 3.**
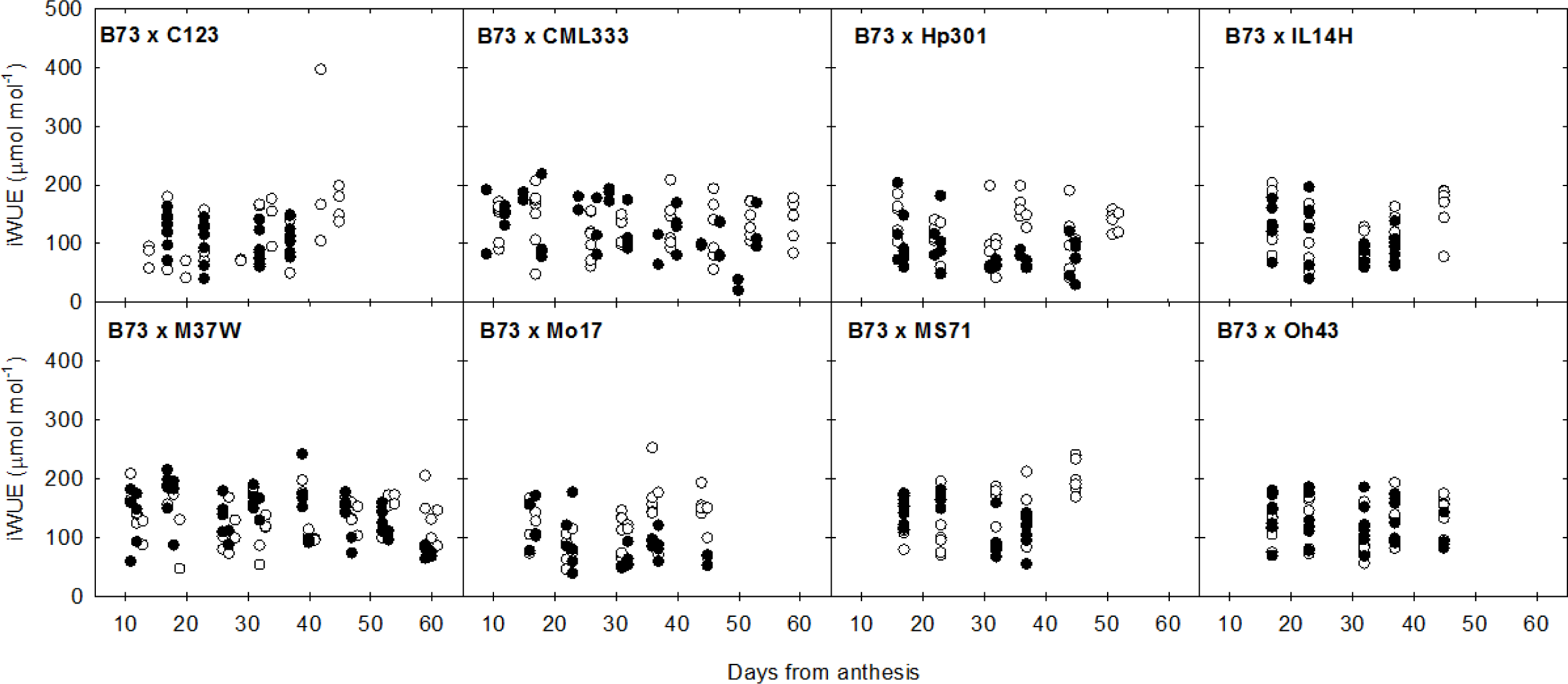
Instantaneous water use efficiency (iWUE = *A*/g_s_) of the leaf subtending the ear measured during grain filling in 2015 for 8 hybrid lines grown at ambient (open circles) and elevated [O_3_] (filled circles).

**Supplemental Table 1.** Parameters describing the quadratic function describing the decline in *A* and g_s_ over time in the leaf subtending the ear, for 10 inbred lines and 8 hybrid lines grown at ambient and elevated [O_3_]. Significant differences in the quadratic fits between ambient and elevated [O_3_] for each inbred or hybrid line are shown in bold font.

| Line | $[O_3]$ | $A$ | | | $g_s$ | | |
| --- | --- | --- | --- | --- | --- | --- | --- |
| | | $y_0$ | $\alpha$ | $\beta$ | $y_0$ | $\alpha$ | $\beta$ |
| B73 | Ambient | **25.6** | **0.91** | **-0.027** | 0.16 | 0.0056 | -0.00018 |
|  | Elevated | **35.9** | **-0.15** | **-0.011** | 0.16 | 0.0056 | -0.00018 |
| C123 | Ambient | **10.2** | **1.99** | **-0.045** | **0.026** | **0.015** | **-0.00037** |
|  | Elevated | **49.4** | **-1.99** | **0.020** | **0.37** | **-0.014** | **0.00013** |
| CML333 | Ambient | 30.8 | 0.22 | -0.014 | 0.19 | 0.00078 | -0.00007 |
|  | Elevated | 30.8 | 0.22 | -0.014 | 0.19 | 0.00078 | -0.00007 |
| HP301 | Ambient | **24.5** | **0.80** | **-0.026** | 0.14 | 0.0052 | -0.00017 |
|  | Elevated | **29.6** | **0.13** | **-0.015** | 0.14 | 0.0052 | -0.00017 |
| IL14H | Ambient | **61.5** | **-1.89** | **0.011** | 0.59 | -0.025 | 0.00025 |
|  | Elevated | **63.4** | **-2.62** | **0.026** | 0.59 | -0.025 | 0.00025 |
| M37W | Ambient | 22.3 | 0.0077 | -0.0046 | 0.13 | 0.00087 | -0.00004 |
|  | Elevated | 22.3 | 0.0077 | -0.0046 | 0.13 | 0.00087 | -0.00004 |
| Mo17 | Ambient | **25.9** | **0.30** | **-0.015** | 0.073 | 0.0064 | -0.00016 |
|  | Elevated | **16.6** | **0.75** | **-0.022** | 0.073 | 0.0064 | -0.00016 |
| MS71 | Ambient | **48.6** | **-1.19** | **0.0035** | **0.37** | **-0.011** | **0.000066** |
|  | Elevated | **60.4** | **-2.47** | **0.025** | **0.42** | **-0.018** | **0.00020** |
| NC338 | Ambient | **24.8** | **0.28** | **-0.011** | **0.096** | **0.0046** | **-0.00012** |
|  | Elevated | **30.6** | **0.012** | **-0.010** | **0.047** | **0.0084** | **-0.00019** |
| OH43 | Ambient | **20.0** | **0.97** | **-0.022** | 0.17 | 0.0025 | -0.00009 |
|  | Elevated | **28.8** | **0.28** | **-0.014** | 0.17 | 0.0025 | -0.00009 |
| B73 x C123 | Ambient | **4.72** | **1.28** | **-0.026** | **0.18** | **0.0071** | **-0.00021** |
|  | Elevated | **17.0** | **0.10** | **-0.0089** | **0.099** | **0.0066** | **-0.00017** |
| B73 x CML333 | Ambient | **15.7** | **0.41** | **-0.0078** | 0.11 | 0.0051 | -0.00010 |
|  | Elevated | **16.7** | **0.34** | **-0.0097** | 0.11 | 0.0051 | -0.00010 |
| B73 x HP301 | Ambient | **6.35** | **0.93** | **-0.017** | 0.0063 | 0.014 | -0.00024 |
|  | Elevated | **15.5** | **0.19** | **-0.0082** | 0.0063 | 0.014 | -0.00024 |
| B73 x IL14H | Ambient | **7.52** | **0.98** | **-0.020** | 0.041 | 0.012 | -0.00025 |
|  | Elevated | **24.0** | **-0.22** | **-0.0054** | 0.041 | 0.012 | -0.00025 |
| B73 x M37W | Ambient | **15.4** | **0.19** | **-0.0038** | 0.13 | 0.0022 | -0.00005 |
|  | Elevated | **16.8** | **0.28** | **-0.0083** | 0.13 | 0.0022 | -0.00005 |
| B73 x Mo17 | Ambient | **3.37** | **1.27** | **-0.025** | 0.075 | 0.010 | -0.00023 |
|  | Elevated | **24.1** | **-0.41** | **-0.0014** | 0.075 | 0.010 | -0.00023 |
| B73 x MS71 | Ambient | **1.02** | **1.42** | **-0.028** | **0.033** | **0.0116** | **-0.00024** |
|  | Elevated | **21.4** | **-0.28** | **-0.0031** | **0.071** | **0.0037** | **-0.00010** |
| B73 x OH43 | Ambient | **-4.85** | **1.71** | **-0.030** | **-0.15** | **0.023** | **-0.00039** |
|  | Elevated | **24.4** | **-0.19** | **-0.0053** | **0.17** | **0.00060** | **-0.00007** |

## REFERENCES

Ahmadzadeh A., Lee E.A. & Tollenaar M. (2004) Heterosis for leaf CO_2_ exchange rate during the grain-filling period in maize. Crop Science 44, 2095–2100.

Ainsworth E.A. (2017) Understanding and improving global crop response to ozone. Plant Journal, DOI: 10.1111/tpj.13298.

Ainsworth E.A., Yendrek C.R., Sitch S., Collins W.J. & Emberson L.D. (2012) The effects of tropospheric ozone on net primary production and implications for climate change. Annual Review of Plant Biology 63, 637–661.

Albergoni F.G., Basso B., Pe E. & Ottaviano E. (1983) Photosynthetic rate in maize. Inheritance and correlation with morphological traits. Maydica 28, 439–448.

Avnery S., Mauzerall D.L., Liu J. & Horowitz L.W. (2011) Global crop yield reductions due to surface ozone exposure: 1. Year 2000 crop production losses and economic damage. Atmospheric Environment 45, 2284–2296.

Bagard M., Jolivet Y., Hasenfratz-Sauder M-P., Gérard J., Dizengremel P. & Le Thiec D. (2015) Ozone exposure and flux-based response functions for photosynthetic traits in wheat, maize and poplar. Environmental Pollution 206, 411–420.

Bänziger M., Edmeades G.O. & Lafitte H.R. (1999) Selection for drought tolerance increases maize yields across a range of nitrogen levels. Crop Science 39, 1035–1040.

Borrás L., Slafer G.A. & Otegui M.E. (2004) Seed dry weight response to source-sink manipulations in wheat, maize and soybean: a quantitative reappraisal. Field Crops Research 86, 131–146.

Christensen J.H., Hewitson B., Busuioc A., Chen A., Gao X., Held I.,…, Whetton P. (2007) Regional climate projections. In Climate Change 2007: The Physical Science Basis. Contribution of Working Group I to the Fourth Assessment Report of the Intergovernmental Panel on Climate Change (eds S. Solomon, D. Qin, M. Manning, Z. Chen, M. Marquis, K.B. Averyt, M. Tignor, H.L. Miller), pp. 847–940. Cambridge University Press, Cambridge, UK, & New York, NY.

Cook B. I., Smerdon J. E., Seager R. & Coats S. (2014) Global warming and 21^st^ century drying. Climate Dynamics 43, 2607–2627.

Ding L., Wang K.J., Jiang G.M., Biswas D.K., Hu X., Li F. & Li Y.H. (2005) Effects of nitrogen deficiency on photosynthetic traits of maize hybrids released in different years. Annals of Botany 96, 925–930.

Emberson L.D., Pleijel H., Ainsworth E.A., van den Berg M., Ren W., Osborne S.,…, Van Dingenen R. (2017) Ozone effects on crops and consideration in crop models. European Journal of Agronomy, accepted.

Farquhar G.D. & Sharkey T.D. (1982) Stomatal conductance and photosynthesis. Annual Review of Plant Physiology 33, 317–345.

Felzer B.S., Cronin T., Reilly J.M., Melillo J.M. & Wang X. (2007) Impacts of ozone on trees and crops. C.R. Geoscience 339, 784–798.

Feng Z., Pang J., Kobayashi K., Zhu J. & Ort D. R. (2011) Differential responses in two varieties of winter wheat to elevated ozone concentration under fully open-air field conditions, Global Change Biology 17, 580–591.

Feng Z., Wang L., Noichi I., Kobayashi K., Yamakawa T. & Zhu J. (2010) Apoplastic ascorbate contributes to the differential ozone sensitivity in two varieties of winter wheat under fully open-air field conditions. Environmental Pollution 158, 3539–3545.

Feng Z., Wang L., Pleijel H., Zhu J. & Kobayashi K. (2016) Differential effects of ozone on photosynthesis of winter wheat among cultivars depend on antioxidative enzymes rather than stomatal conductance. Science of the Total Environment 572, 404–411.

Fiscus E.L., Booker F.L. & Burkey K.O. (2005) Crop responses to ozone: uptake, modes of action, carbon assimilation and partitioning. Plant Cell and Environment 28, 997–1011.

Flint-Garcia S.A., Thuillet A.C., Yu J., Pressoir G., Romero S.M., Mitchell S.E., Doebley J., Kresovich S., Goodman M.M. & Buckler E.S. (2005) Maize association population: a high-resolution platform for quantitative trait locus dissection. Plant Journal 44, 1054–1064.

Gillespie K.M., Xu F., Richter K.T., McGrath J.M., Markelz R.J.C., Ort D.R., Leakey A.D.B. & Ainsworth E.A. (2012) Greater antioxidant and respiratory metabolism in field-grown soybean exposed to elevated O_3_ and two CO_2_ concentrations. Plant Cell & Environment 35, 169–184.

Heath R.L. (1988) Biochemical mechanisms of pollutant stress. In Heck WW, Taylor OC, Tingey DT, eds, Assessment of Crop Losses from Air Pollutants. Elsevier, New York, pp 259–286.

Heck W.W., Adams R.M., Cure W.W., Heagle A.S., Heggestad H.E., Kohut R.J., Kress L.W., Rawlings J.O. & Taylor O.C. (1983) A reassessment of crop loss from ozone. Environmental Science & Technology 17, 572A–581A.

Keller T. & Häsler R., 1984. The influence of a fall fumigation with ozone on the stomatal behavior of spruce and fir. Oecologia 64, 284–286.

Kress L.W. & Miller J.E. (1985) Impact of ozone on field-corn yield. Canadian Journal of Botany 63, 2408–2415.

Krupa S.V. & Manning W.J. (1988) Atmospheric ozone: formation and effects on vegetation. Environmental Pollution 50, 101–137.

Leakey A.D.B., Uribelarrea M., Ainsworth E.A., Naidu S. L, Rogers A., Ort D.R. & Long S.P. (2006) Photosynthesis, productivity, and yield of maize are not affected by open-air elevated of CO_2_ concentration in the absence of drought. Plant Physiology 140, 779–790.

Leitao L., Bethenod O. & Biolley J.P. (2007) The impact of ozone on juvenile maize (*Zea mays* L.) plant photosynthesis: effects on vegetative biomass, pigmentation, and carboxylases (PEPc and Rubisco). Plant Biology 9, 478–488.

Leitao L., Maoret J.J. & Biolley J.P. (2007b) Changes in PEP carboxylase, rubisco and rubisco activase mRNA levels from maize (*Zea mays*) exposed to a chronic ozone stress. Biological Research 40, 137–153.

Lin M., Horowitz L.W., Payton R., Fiore A.M. & Tonnesen G. (2017) US surface ozone trends and extremes from 1980 to 2014: quantifying the roles of rising Asian emissions, domestic controls, wildfires, and climate. Atmospheric Chemistry & Physics 17, 2943–2970.

Lombardozzi D., Sparks J.P., Bonan G. & Levis S. (2012) Ozone exposure causes a decoupling of conductance and photosynthesis: implications for the Ball-Berry stomatal conductance model. Oecologia 169, 651–659.

Markelz R.J.C., Strellner R.S. & Leakey A.D.B. (2011) Impairment of C4 photosynthesis by drought is exacerbated by limiting nitrogen and ameliorated by elevated [CO_2_] in maize. Journal of Experimental Botany 62, 3235–3246.

McGrath J.M., Betzelberger A.M., Wang S., Shook E., Zhu X-G., Long S.P. & Ainsworth E.A. (2015) An analysis of ozone damage to historical maize and soybean yields in the United States. Proceedings of the National Academy of Science USA 112, 14390–14395.

Mills G., Buse A., Gimeno B., Bermejo V., Holland M., Emberson L. & Pleijel H. (2007) A synthesis of AOT40-based response functions and critical levels of ozone for agricultural and horticultural crops. Atmospheric Environment 41, 2630–2643.

Mulchi C., Rudorff B., Lee E., Rowland R. & Pausch R. (1995) Morphological responses among crop species to full-season exposures to enhanced concentrations of atmospheric CO_2_ and O_3_. Water, Air and Soil Pollution 85, 1379–1386.

Neiff N, Trachsel S, Valentinuz OR, Balbi CN & Andrade FH (2016) High temperatures around flowering in maize: effects on photosynthesis and grain yield in three genotypes. Crop Science 56, 2702–2712.

Pang J., Kobayashi K. & Zhu J. (2009) Yield and photosynthetic characteristics of flag leaves in Chinese rice (*Oryza sativa* L.) varieties subjected to free-air release of ozone. Agriculture, Ecosystems & Environment 132, 203–211.

Paoletti E. & Grulke N.E. (2005) Does living in elevated CO_2_ ameliorate tree response to ozone? A review on stomatal responses. Environmental Pollution 137, 483–93.

Paoletti E. & Grulke N.E. (2010) Ozone exposure and stomatal sluggishness in different plant physiognomic classes. Environmental Pollution 148, 2664–2671.

Pinder JE, Wiener JG, Smith MH (1978) The Weibull distribution: a new method of summarizing survivorship data. Ecology 59, 175–179.

Pino ME, Mudd JB, Bailey-Serres J (1995) Ozone-induced alterations in the accumulation of newly synthesized proteins in leaves of maize. Plant Physiology 108, 777–785

Reich P.B. (1984) Loss of stomatal function in ageing hybrid poplar leaves. Annals of Botany 53, 691–698.

Reich P.B. (1987) Quantifying plant response to ozone: a unifying theory. Tree Physiology 3, 63–91.

Rudorff B.F.T., Mulchi C.L., Lee E.H., Rowland R. & Pausch R. (1996) Effects of enhanced O_3_ and CO_2_ enrichment on plant characteristics in wheat and corn. Environmental Pollution 94, 553–60.

Sitch S., Cox P., Collins W.J. & Huntingford C. (2007) Indirect radiative forcing of climate change through ozone effects on the land-carbon sink. Nature 448, 791–794.

Thomas H. & Howarth C.J. (2000) Five ways to stay green. Journal of Experimental Botany 51, 329–337.

Tjoelker M.G., Volin J.C., Oleksyn J. & Reich P.B. (1995) Interaction of ozone pollution and light effects on photosynthesis in a forest canopy experiment. Plant, Cell & Environment 18, 895–905.

Tollenaar M., Ahmadzadeh A. & Lee E.A. (2004) Physiological basis of heterosis for grain yield in maize. Crop Science 44, 2086–2094.

Valentinuz O.R. & Tollenaar M. (2004) Vertical profile of leaf senescence during the grain-filling period in older and newer maize hybrids. Crop Science 44, 827–834.

VanLoocke A., Betzelberger A.M., Ainsworth E.A. & Bernacchi C.J. (2012) Rising ozone concentrations decrease soybean evapotranspiration and water use efficiency whilst increasing canopy temperature. New Phytologist 195, 164–171.

von Caemmerer S. (2000) Biochemcial models of leaf photosynthesis. CSIRO Publishing, Collingwood, Australia.

Wagg S., Mills G., Hayes F., Wilkinson S., & Davies W.J. (2013) Stomata are less responsive to environmental stimuli in high background ozone in *Dactylis glomerata* and *Ranunculus acris*. Environmental Pollution 175, 82–91.

Walsh J, Wuebbles D, Hayhoe K, Kossin J, Kunkel K, Stephens G, Thorne P, Vose R, Wehner M, Willis J, Anderson D, Doney S, Feely R, Hennon P, Kharin V, Knutson T, Landerer F, Lenton T, Kennedy J, Somerville R (2014) Ch. 2: Our Changing Climate. Climate Change Impacts in the United States: The Third National Climate Assessment., Melillo JM, Richmond TC, Yohe GW, Eds., U.S. Global Change Research Program, 19–67.

Wilkinson S. & Davies W.J. (2009) Ozone suppresses soil drying- and abscisic acid (ABA)- induced stomatal closure via an ethylene-dependent mechanism. Plant Cell & Environment 32, 949–959.

Wittig V.E., Ainsworth E.A. & Long S.P (2007) To what extent to current and projected increases in surface ozone affect photosynthesis and stomatal conductance of trees? A meta-analytic review of the last 3 decades of experiments. Plant Cell & Environment 30, 1150–1162.

Wolfe D.W., Henderson D.W., Hsiao T.C. & Alvino A. (1988) Interactive water and nitrogen effects on senescence of maize. II. Photosynthetic decline and longevity of individual leaves. Agronomy Journal 80, 865–870.

Yendrek C.R., Tomaz T., Montes C.M., Cao Y., Morse A.M., Brown P.J., McIntyre L.M., Leakey A.D.B. & Ainsworth E.A. (2017) High-throughput phenotyping of maize leaf physiological and biochemical traits using hyperspectral reflectance. Plant Physiology 173, 614–626.

Yu J., Holland J.B., McMullen M.D. & Buckler E.S. (2008) Genetic design and statistical power of nested association mapping in maize. Genetics 178, 539–551.

Zheng Y., Shimizu H. & Barnes J.D. (2002) Limitations to CO_2_ assimilation in ozone-exposed leaves of Plantago major. New Phytologist 155, 67–78.

